# Virome-mediated ecological firewalls govern pathogen eradication in the gut microbiome

**DOI:** 10.64898/2026.06.02.729649

**Authors:** Andrea Tabi, Ricard Solé

## Abstract

Microbiome recovery after antibiotic-induced dysbiosis is often unpredictable, and the ecological mechanisms governing successful gut microbiome restoration remain unclear. Here we introduce a novel dynamical model of bacteria-phage-pathogen interactions to examine how gut virome reshapes gut microbiome states. The model identifies pathogen eradication thresholds and shows that recovery is strongly history-dependent, with bistability allowing either clearance or persistence depending on past conditions. Viral adsorption rates strongly regulate invasion probability and recurrence susceptibility, while microbial diversity enhances stability. Furthermore, we identify an ecological “firewall” mechanism in which the resident microbiome maintains a standing virome community that suppresses pathogen invasion and stabilizes healthy microbiome. These results reveal general ecological principles governing virus-mediated microbiome recovery and inform the design of virome-based therapeutic strategies.

## I. INTRODUCTION

The gut microbiome (GM) is a complex ecosystem that, in many respects, behaves as a species-rich ecological community [13], with interacting populations competing for resources, exchanging metabolites, and responding to environmental perturbations [31]. Its structure and dynamics emerge from multiscale processes ranging from microbial metabolism and interspecies interactions to host-mediated constraints, placing the GM at the intersection of ecology, evolution, and physiology. Perhaps not surprisingly, many approaches to the dynamics of the GM are grounded in ecological and evolutionary concepts, including stability [52], alternative stable states [16, 29, 30], succession [11, 18], resilience [12], and metapopulation structure [27, 28]. At the same time, the GM exhibits distinctive features that distinguish it from most free-living ecosystems, such as high levels of functional redundancy in taxa, rapid evolutionary turnover, and strong coupling to host regulatory mechanisms, including immune and neuroendocrine pathways [10, 36]. These idiosyncrasies complicate direct ecological analogies while motivating the development of new mathematical and computational models [2, 4, 8, 9, 20, 22, 47, 51, 53]. At the same time, the GM has been described as an “ecosystem on a leash,” emphasizing the role of host control in shaping microbial dynamics and constraining evolutionary outcomes [15]. Health benefits associated with the microbiome thus emerge from a delicate interplay between microbial interactions and host-mediated selection and regulation. As a result, GM has become a central focus of contemporary biomedical research, with a rapidly growing body of evidence linking its composition and dynamics to a wide range of health outcomes, including metabolic disorders, immune dysfunction, neurological conditions, and susceptibility to infection [6, 10, 39, 41]. This recognition has, in turn, motivated the exploration of microbiome-targeted interventions, including antibiotics [14] dietary modulation and probiotics [25], fecal microbiota transplantation [43] and, more recently, virome-based therapies [54].

More specifically, antibiotic-mediated damage of the gut microbiome opens an opportunistic niche for pathogenic colonisers such as drug-resistant *Enterobacteriaceae, Clostridioides difficile*, vancomycin-resistant *Enterococcus* and *Salmonella* spp. [14]. Antibiotics acutely and persistently perturb the gut microbial community by impacting microbial community composition and structure and eroding important taxa such as Bacteroidetes, Firmicutes and Actinobacteria. As a remedy for antibiotic-induced infections, recent advances across clinical microbiome research have shown the high efficacy of fecal microbiota transplantation (FMT). Synthesized clinical literature report that FMT achieves 80-90% success rates in *C. difficile* infections (CDI) and has an emerging roles in other metabolic, autoimmune, and neurological disorders [21, 37]. Several studies demonstrated that FMT restores microbial diversity, changes in metabolic functions, modulates the immune system, affects bacteriophage populations in the gut [24, 32], influences the dynamics of bacterial strains [42]. Despite the high effectiveness of FMT treatments, there are significant drawbacks such as safety concerns and the challenging and expensive donor recruitment and screening process [53]. Therefore, future FMT research requires more focus on host immune status and non-bacterial components [24].

Recent works suggest that fecal virome transplantation (FVT) have been similarly effective in treating rCDI, possibly through bacteriophage-mediated modulation of the gut microbiome [7, 40]. This method might potentially reduce risks of infections following FVT treatment by removing harmful eukaryotic RNA viruses and increase donor suitability [40]. Bacteriophages have a large impact on different ecosystems from the oceans to the human gut [3, 33, 49]. The adaptive capacity of phages plays a crucial role in multidrug resistance and protecting the gut microflora, and preserving its functional robustness during antibiotic stress [35, 45]. Despite growing evidence that bacteriophages are major regulators of microbial communities, most theoretical models of the gut microbiome focus exclusively on bacterial interactions and neglect viral dynamics. Incorporating the virome is therefore essential for understanding microbiome stability and pathogen invasion. However, the exact mechanisms of how phages mediate and restore bacterial community structure are largely unknown. Bacteriophages can modify intrinsic growth rates and interaction coefficients of bacterial species, e.g. via horizontal gene transfer of beneficial metabolic genes [19].

In this work we identify ecological virus-mediated mechanisms driving microbiome restoration. First, we derive eradication thresholds that determine when pathogen growth can be suppressed by bacterial competition and viral adsorption. Second, we identify alternative stable states, i.e. path-dependent recovery dynamics in which invasion probability and recurrence susceptibility depend on initial microbiome composition. Third, we identify a virome-mediated “ecological firewall” mechanism whereby resident microbial communities maintain viral populations that suppress pathogen invasion. Finally, we show that increasing virome diversity strengthen invasion resistance.

## II. METHODS

To investigate the mechanisms by which virome transplantation can suppress pathogens and promote recovery from dysbiosis, we develop a minimal but mechanistically explicit dynamical model of a gut microbial ecosystem composed of commensal bacteria, bacteriophages, and a pathogenic population. Figure 1a-b summarizes the ecological scenario motivating the model, illustrating how antibiotic perturbation can disrupt the resident microbiome, allowing *C. difficile* to expand and drive the system into a dysbiotic state, and how subsequent virome transplantation can restore microbial balance. At the conceptual level (Fig. 1b), the system consists of three interacting compartments: commensal bacterial populations {*B*_*i*_} (*i* = 1, …, *N* ), a pathogen *C*, and a virome {*P*_*j*_} (*j* = 1, …, *M* ). The modelling framework is designed to capture the essential ecological processes operating across these compartments, including intrinsic microbial growth under resource limitation, competitive and facilitative interactions among bacterial taxa, phage-host dynamics, and the targeted introduction of exogenous phages during virome transplantation. These processes are formalized in the mathematical model, which follows a generalized Lotka-Volterra (gLV) approach. Our system of *N* + *M* + 1 equations, shown in Fig.1c.

**Figure 1.**
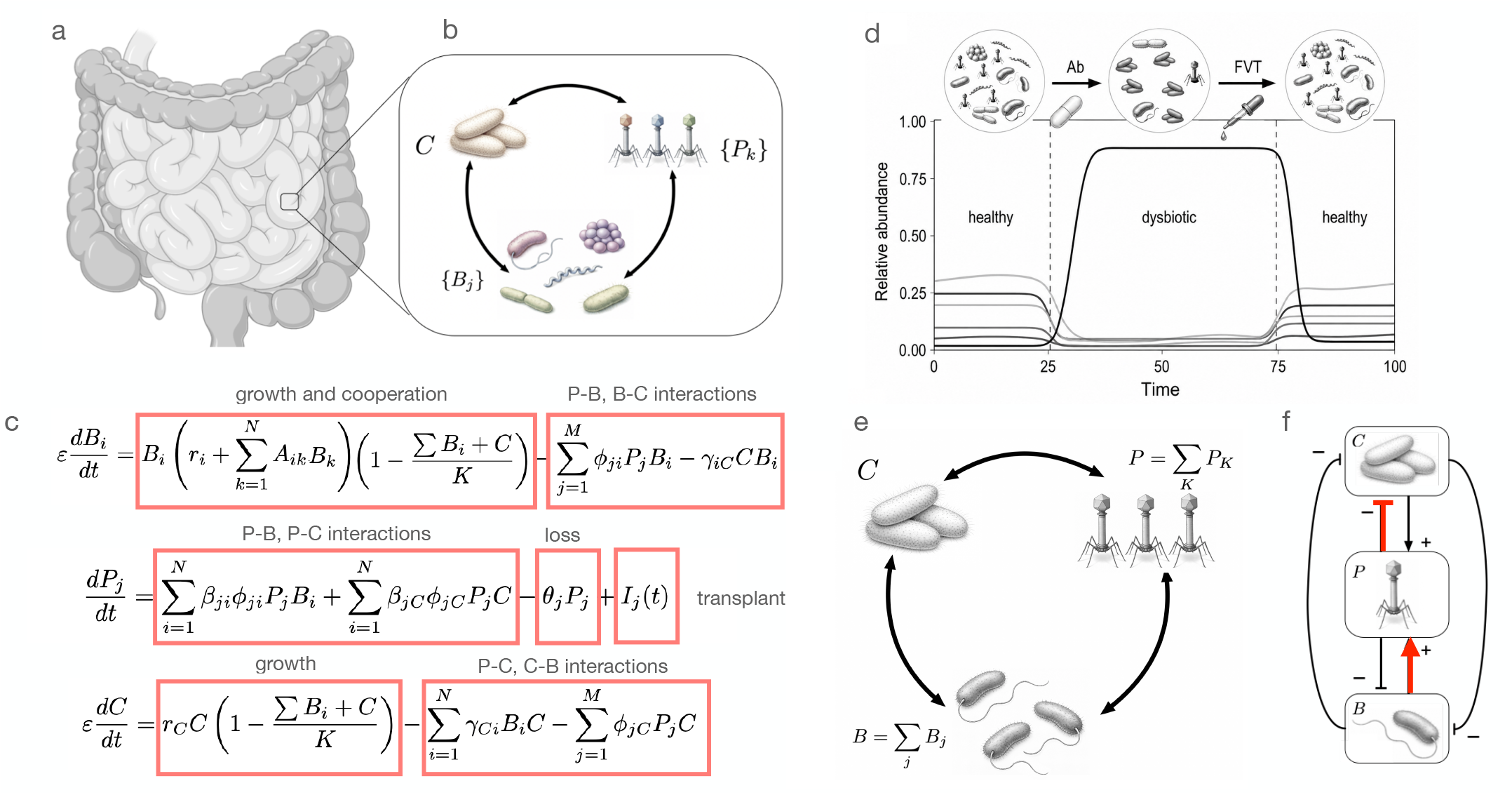
Ecological and mathematical framework for fecal virome transplantation in the gut microbiome. (a) The gut ecosystem is represented as a community of interacting microbial populations inhabiting the intestinal environment. (b) Conceptual interaction network underlying the model. Commensal bacterial taxa {*B*_*j*_}, an opportunistic pathogen population *C*, and bacteriophages {*P*_*k*_} form a coupled ecological system. Phages are sustained by bacterial hosts and exert top–down control on both commensal and pathogenic populations, generating feedbacks that shape community composition and ecosystem stability. (c) Mean-field dynamical model describing the coupled bacteria–phage system. Commensal bacteria grow logistically, with microbial facilitation, and are subject to phage predation. The pathogen population *C* competes with resident bacteria and is likewise targeted by phages. Phage populations increase through infection of bacterial hosts and decline through loss processes; the term *I*_*j*_ (*t*) represents the external input associated with virome transplantation. Together, these equations define the ecological dynamics governing transitions between healthy and dysbiotic states. (d) Representative dynamical trajectory illustrating microbiome disruption and recovery. Antibiotic treatment (Ab) drives the system from a healthy state into a dysbiotic regime characterized by pathogen domination and loss of microbial diversity. Subsequent fecal virome transplantation (FVT) restores phage-mediated ecological control, suppresses pathogen overgrowth, and returns the system to a healthy state. (e) Coarse-grained representation of the model. The full microbial community is reduced to three interacting compartments: commensal bacteria (*B* = ∑ _*j*_ *B*_*j*_), pathogen (*C*), and virome (*P* = ∑ _*k*_ *P*_*k*_). This reduced framework captures the essential ecological feedbacks responsible for pathogen suppression, microbiome resilience, and recovery following virome transplantation. The architecture of the interactions is summarized in (f) where direct and indirect effects are displayed, along with two main interactions (in red) responsible for the maintenance of the healthy state.

Describing the coupled dynamics between the microbiome *{B*_*k*_*}*, the virome *{P*_*j*_*}* and the pathogen population *C*. Here 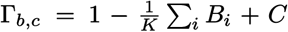 introduces the competition for space between the resident microbiome and the *C. difficile* population. In this formulation, commensal bacteria and the pathogen undergo logistic growth constrained by a shared carrying capacity, reflecting global resource limitation in the gut environment, while bottom-up microbial dynamics are coupled to top-down control mediated by bacteriophages that infect both commensals and the pathogen with distinct efficiencies. The virome is therefore treated as an active regulatory layer capable of reshaping community structure and stability. This community-level defense mechanism can be interpreted as an ecological firewall, by which the resident microbiome indirectly protects itself by maintaining a standing virome, which then suppresses pathogen invasion. Rather than aiming for taxonomic realism, the model focuses on functional interactions that govern ecosystem-level transitions between healthy and dysbiotic states.

## III. RESULTS

### A. Eradication thresholds

To understand the conditions under which the pathogen can be eliminated from the system, we first derive eradication thresholds from our coarse-grained model (see SM for details). Such thresholds identify the parameter regimes in which pathogen growth can no longer compensate for suppressive ecological interactions. In an ecological and epidemiological context, threshold conditions provide a useful analytical tool to define transitions between persistence and extinction [1, 46]. In our context, they allow us to quantify how competitive pressure from the resident microbiome and additional control from bacteriophages limit the growth of *C. difficile*. By first analyzing a bacteria–pathogen system without phages and then extending the analysis to the virus-mediated model, we can isolate the mechanisms that determine when pathogen eradication becomes possible.

First, we estimate the threshold conditions for pathogen clearance (*C* = 0) in a bacteria-pathogen model. The original set of equations (Fig.1c) is thus simplified to a set of *N* + 1 equations, and a further simplification is obtained by using a coarse-grained Bacterium-*C*.*difficile* model (see SM for details). Specifically, if we assume a homogeneous set of parameters (i. e. all *B*_*k*_ exhibit the same growth, death, and interaction rates), the system reduces to a two-dimensional model, where *B* = ∑_*k*_*B*_*k*_ is the aggregated microbiome population as described by equations (1-3) (see Materials and Methods and SM).

A starting pont of our analysis involves a limit case where no virome exists (*P*_*i*_ = 0, *i* = 1, …, *M* ; Fig. 2a). This results in a competition model that can be easily analysed using the coarse-grained approximation. The boundary equilibria of the no-virome system are *E*_0_ = (0, 0), *E*_*H*_ = (*K*, 0), and *E*_*D*_ = (0, *K*) (see Fig.2c). The origin is an unstable source. The healthy equilibrium *E*_*H*_ = (*K*, 0), which corresponds to pathogen clearance, has negative eigenvalues *λ*_*B*_ = − (*r*_*B*_ + *αK*) and *λ*_*C*_ = − *γ*_*CB*_*K*, so it is stable. Similarly, the dysbiotic equilibrium *E*_*D*_ = (0, *K*), corresponding to pathogen persistence, has eigenvalues *λ*_*B*_ = − *γ*_*BC*_*K* and *λ*_*C*_ = − *r*_*C*_, and is also stable. .

**Figure 2.**
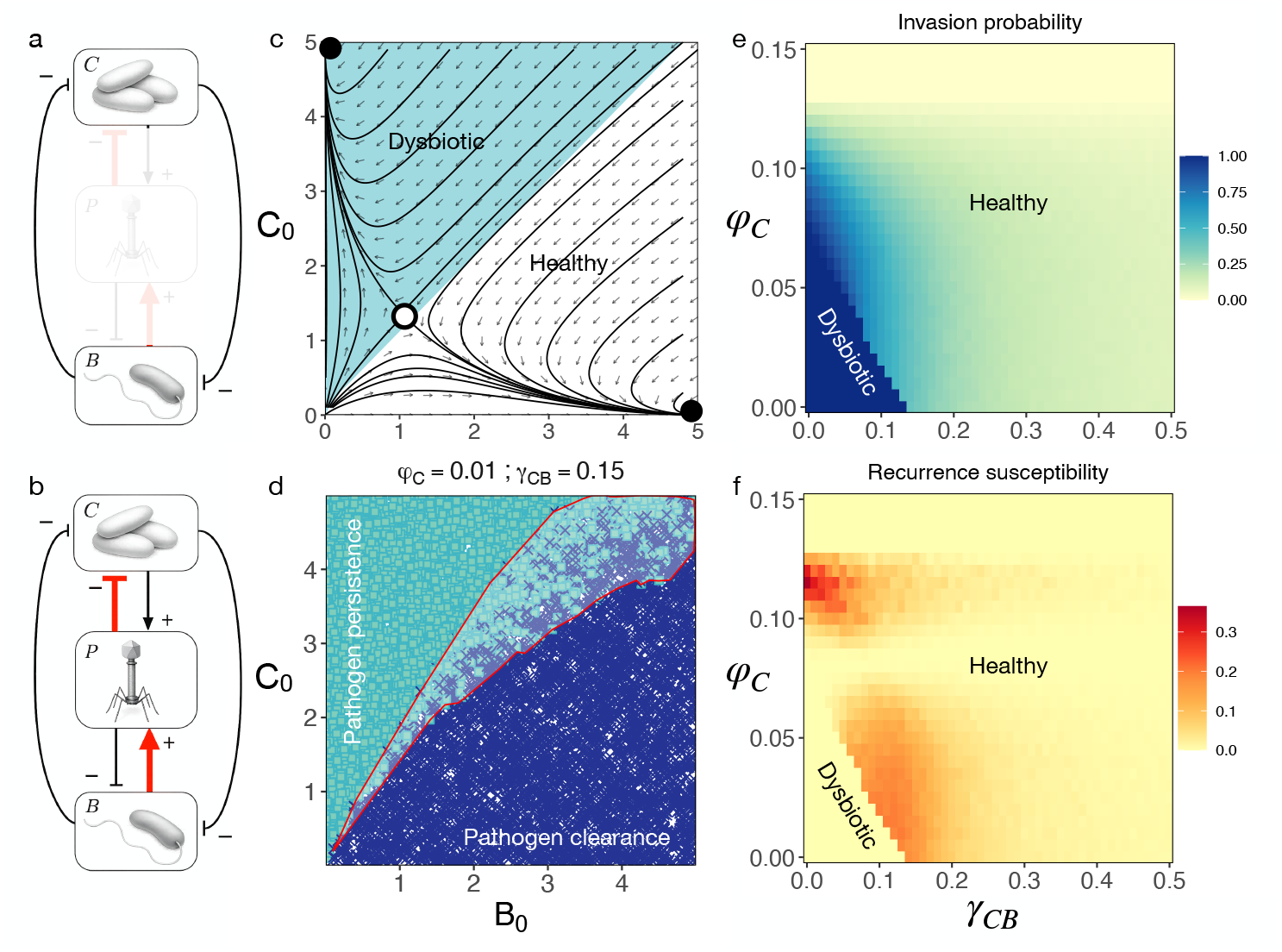
Healthy-dysbiotic transitions under coarse-grained models. (a) In the absence of a virome, the resident microbiome is represented by the aggregate abundance (*B* = ∑_*i*_ *B*_*i*_), which competes with the pathogen population (C) for shared ecological resources. (c) shows the phase portrait of this reduced coarse-grained system. Stable and unstable states are indicated by empty and filled circles, respectively. The system exhibits competitive exclusion, with two alternative outcomes: pathogen clearance (healthy state, *C*^∗^ = 0, dashed areas) and pathogen persistence (dysbiotic state, *C*^∗^ *>* 0). (b) Coarse-grained virus-mediated model including phage interactions between the resident bacteria and the pathogen. (d) Basin of attraction in the Bacterium–Phage-*C. difficile* model for viral adsorption rates *ϕ*_*C*_ = 0.01 and bacterial suppression *γ*_*CB*_ = 0.15. Points show initial conditions (*B*_0_, *C*_0_) and are colored by the long-term equilibrium outcome *C*^∗^, illustrating that *C. difficile* invasion depends on initial microbiome state. The red outlined region represents the recurrence-susceptible region, where both clearance (*C*^∗^ = 0) and persistence (*C*^∗^ *>* 0) are possible (i.e., alternative outcomes occur depending on initial conditions). (e) *C. difficile* invasion probability across *γ*_*cb*_ and *ϕ*_*C*_ computed as the fraction of initial conditions leading to persistence (*C*^∗^ *>* 0). Recurrence susceptibility across *γ*_*CB*_ and *ϕ*_*C*_ computed as the fraction of initial conditions compatible with both clearance and persistence.

How is the complete model affected by the pathogen, as described in terms of an invasion process? In its absence (*C* = 0), the CGM gives an equilibrium value for the microbiome *B*_*H*_ = *θ/*(*β*_*B*_*ϕ*_*B*_), and a virome population:

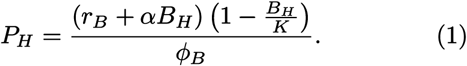

For *C* ≪ 1, the pathogen equation in Eq. (20) can be linearized around the healthy state. Keeping only terms proportional to *C* gives

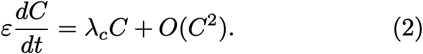

where the pathogen invasion exponent is

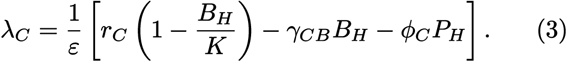

The pathogen invades when *λ*_*C*_ *>* 0 while the healthy state excludes the pathogen when *λ*_*C*_ *<* 0. The transition between pathogen exclusion and pathogen invasion occurs at the critical boundary defined by (see SM):

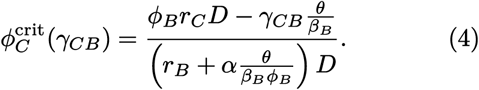

where we define *D* = 1 − *θ/*(*β*_*B*_*ϕ*_*B*_*K*). For *D >* 0, the critical boundary can be written as a linear relation:

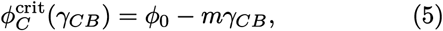

where *ϕ*_0_ is a constant.

Antibiotic treatment primarily targets the resident microbiome, causing a rapid decline in bacterial biomass. At the coarse-grained level this can be represented by adding an external mortality term,

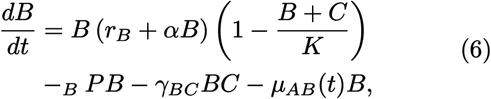

where *µ*_*AB*_(*t*) describes a time-dependent mortality rate due to antibiotic exposure. Immediately after treatment, the resident biomass drops from *B*^∗^ to a lower value *B*_*A*_. Because phage persistence depends on bacterial hosts, the virome is also indirectly affected and decreases from *P*^∗^ to *P*_*A*_. The pathogen growth rate then becomes

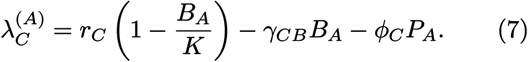

Several effects now act in the same direction. First, reduction of *B*_*A*_ decreases competitive exclusion and antagonistic suppression. Second, lower bacterial density weakens maintenance of the resident virome, reducing phage-mediated pathogen control. Third, the pathogen experiences reduced competition for resources. As a consequence, a system initially satisfying *λ*_*C*_ *<* 0 can be shifted into a dysbiotic regime characterized by 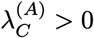.

The pathogen then expands exponentially during the early stages of invasion, following

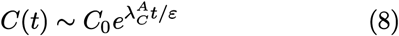

In this picture, dysbiosis is not simply the result of pathogen introduction. Rather, it emerges because antibiotic perturbation moves the ecosystem across the invasion threshold separating pathogen exclusion from pathogen growth. A virome transplant introduces an external inoculum of bacteriophages (*I*_*F V T*_ ) into the dysbiotic ecosystem. The phage equation becomes

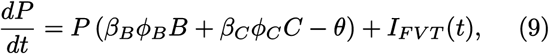

where *I*_*F V T*_ (*t*) represents the virome inoculum. For a pulse-like transplantation at time *t* = *t*_*F*_, i. e. using

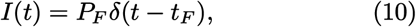

so that the overall phage population contains the resident *P*_*R*_ and transplanted phages *P*_*F*_ :

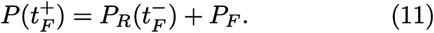

Immediately after transplantation the pathogen invasion eigenvalue becomes

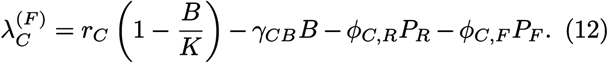

The direct effect of the transplant is therefore to increase the magnitude of the phage suppression term. Pathogen decline occurs whenever 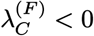. This condition defines a critical therapeutic inoculum

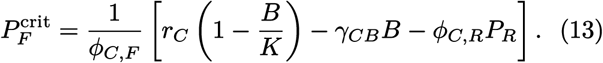

For 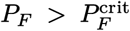, the pathogen growth rate becomes negative and the pathogen population decays. The therapeutic action of VT is not limited to the suppression of direct pathogens. Once the pathogen begins to decline, the resident microbiome experiences reduced competitive pressure and can recover. As *B* increases, two additional feedbacks are activated. First, larger bacterial populations increase niche occupation and antagonistic suppression of the pathogen through the term *γ*_*CB*_*BC*. Second, recovery of the resident microbiome improves phage production by *β*_*B*_*ϕ*_*B*_*BP*, thus strengthening the virome itself.

The healthy state is therefore stabilized by two coupled feedback loops (Fig. 1f), namely:

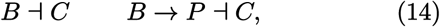

representing direct microbiome-mediated suppression of the pathogen and indirect suppression mediated by bacteriophages, respectively. These feedbacks explain why VT can have long-lasting effects even when the introduced phages themselves are transient. The transplant shifts the system toward a region of state space where the resident microbiome can recover, and once recovery occurs the microbiome sustains the virome that in turn maintains pathogen control. Successful therapy therefore requires more than transient pathogen killing. The intervention must move the system across the separatrix that divides healthy and dysbiotic states. Once this threshold is crossed, ecological feedbacks involving the resident microbiome and the virome drive the system toward recovery. In this interpretation, VT acts as an ecological restoration strategy that re-establishes the coupled bacteria-phage network responsible for maintaining colonization resistance against pathogen invasion.

### B. Path-dependence of pathogen invasion and recurrence

In our coarse-grained Bacterium-Pathogen model, bacteria and the pathogen compete for the same ecological resources. The competition for shared resources (carrying capacity) introduces nonlinear feedbacks leading to alternative stable states corresponding to bacterial dominance (*C*^∗^ = 0) or *C. difficile* persistence (*C*^∗^ *>* 0). Which equilibrium is reached depends on the initial conditions of bacteria (*B*_0_) and *C. difficile* (*C*_0_) densities (Fig.2). Thus, identical parameter values can lead to qualitatively different outcomes depending on initial conditions. The eradication thresholds correspond to the boundary separating the basins of attraction of the pathogen-free and pathogen-persistent equilibria. The observed bistability reflects a switch-like behavior, representing one of the simplest dynamical mechanisms by which biological systems exhibits history-dependence, i.e. memory of prior states. Once the system settles into one attractor, small perturbations do not induce a transition to the other state preserving the outcome of past conditions. This results in path-dependent colonisation. In the virus-mediated model, the system still exhibits alternative stable states (Fig.3a), in which the invasion dynamics of *C*.*difficile* is determined by two factors, the viral adsorption rate (*ϕ*_*C*_) and bacterial suppression rate (*γ*_*CB*_). We computed the invasion probability of *C. difficile* as the fraction of initial conditions leading to persistence across bacterial suppression rates. In the Bacterium-Pathogen model, invasion probability gradually weakens with stronger suppression. While, invasion in the virus-mediated model decreases with bacterial suppression and shows a hard threshold of viral adsorption rate, beyond which invasion of *C*.*difficile* becomes unfeasible (Fig.3b). Recurrence susceptibility, defined as the region of initial conditions compatible with both clearance and persistence outcomes. In the Bacteria-Pathogen model, system dynamics are determined by a single dominant attractor, such that trajectories converge either to pathogen clearance or persistence. As a result, recurrence following perturbations is unlikely. By contrast, in the virus-mediated model recurrence susceptibility becomes substantial and peaks at intermediate values of bacterial suppression and viral adsorption (Fig. 3c). In this regime, virus-mediated pathogen control and bacterial suppression interact to generate bistability, producing mixed-outcome region in the bacteria-pathogen state space. Consequently, small differences in specific regions of initial conditions in bacteria under identical parameter values can result in qualitatively different outcomes. This sensitivity to initial conditions reflects strong influence of virome dynamics and implies an increased likelihood of recurrence following perturbations, as the system can be repeatedly displaced between alternative stable states.

**Figure 3.**
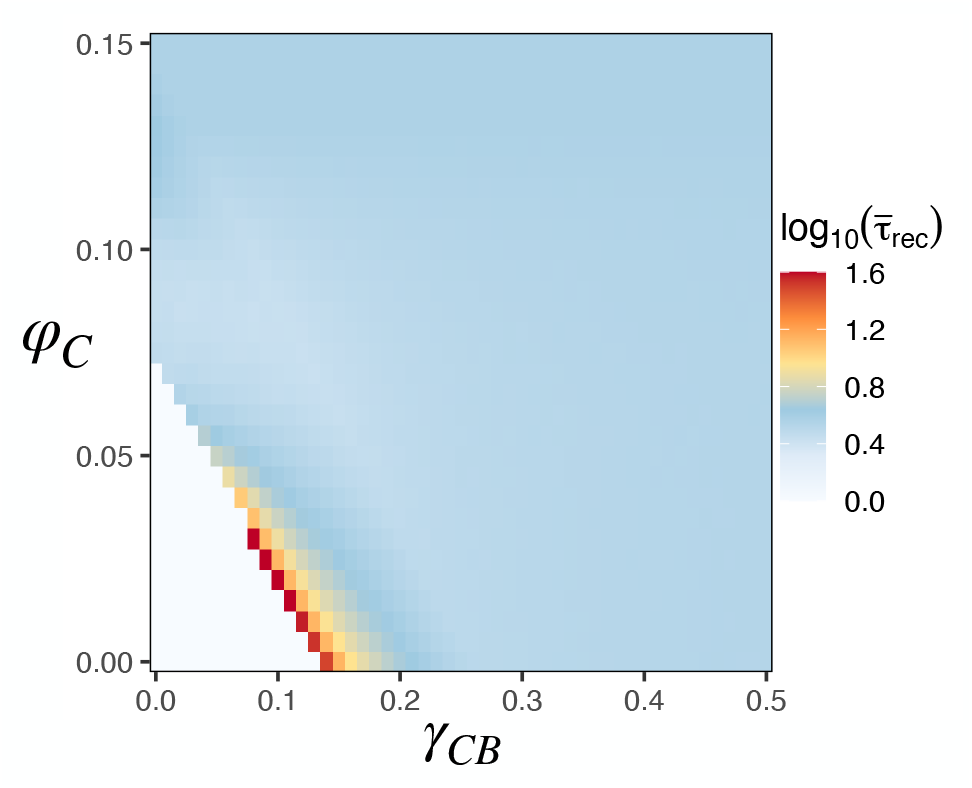
Recovery-time landscape. Mean recovery time 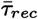, estimated as the inverse dominant eigenvalue of the equilibrium Jacobian, across bacterial suppression (*γ*_*CB*_ ) and viral adsorption (*ϕ*_*C*_ ). Recovery is rapid throughout most of parameter space but becomes markedly slower near the pathogen eradication boundary, where the dominant eigenvalue approaches zero. The resulting ridge of elevated recovery times is consistent with critical slowing down near the transition between pathogen persistence and pathogen clearance.

Importantly, the long-term outcome depends strongly only on the initial composition of the bacterial community and the pathogen, but it is independent of the initial phage abundance. Basin-of-attraction analyses reveal that varying the initial phage densities does not qualitatively alter the system trajectory (Fig. S3 and S4 in SM). Together these results indicate that path dependence arises from the initial microbiome-pathogen balance rather than from the initial viral abundance. The virome influences invasion thresholds through its equilibrium abundance, but the transient initial phage density does not determine the long-term ecological outcome.

### C. Recovery time and transient dynamics

To quantify microbiome resilience, we estimated the local recovery time from the dominant eigenvalue of the Jacobian evaluated at equilibrium,

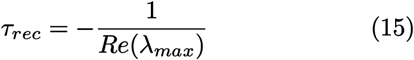

where *λ*_*max*_ denotes the eigenvalue with the largest real part. Larger values correspond to slower recovery following perturbations and therefore reduced ecological resilience. Recovery times were generally short throughout most of parameter space, indicating rapid return toward equilibrium following perturbation (Fig.3). However, a narrow region of elevated recovery times emerges at the boundary, where recovery slows by more than an order of magnitude. This pattern is consistent with critical slowing down near a dynamical transition, where the dominant eigenvalue approaches zero and system resilience weakens.

Overall, transient dynamics dominates recovery including monotonic relaxation and oscillatory transients such as damped oscillations. Dynamical behavior was classified using eigenvalues and trajectories. Systems were first categorized as transient if trajectories converged to equilibrium, i.e. eigenvalues had negative real parts. Among transient dynamics, oscillatory behavior was identified by counting sign changes in deviations from equilibrium *x*(*t*) − *x*^∗^. Trajectories with two or more sign changes were classified as damped oscillations and those with at most one sign change were classified as monotonic relaxation. Almost 86% cases showed oscillatory transients arising from nonlinear feedbacks between bacterial and viral populations. Monotonic relaxation is the second most frequent dynamical behavior (14%) occurring at lower levels of both viral adsorption and bacterial suppression rates. Transitions to sustained oscillations (limit cycles) were identified by eigenvalues with positive real parts and nonzero imaginary part, consistent with Hopf bifurcations.

### D. Diversity effects

The coarse-grained model provides analytical tractability, but it neglects the diversity inherent to real gut ecosystems, where multiple bacterial and phage strains coexist and interact. To assess how diversity influences system dynamics, we extended the framework to multi-strain communities with varying numbers of bacterial and phage populations (Fig.4), introducing small perturbations to their parameter values while keeping the average coefficients to capture functional variability (see for more details in SM).

In the absence of phages, increasing bacterial diversity had no significant effect on *C. difficile* invasion probability (Fig.S6 in SM). Across multi-strain communities, invasion curves largely overlapped, indicating that bacterial diversity alone does not substantially alter invasion outcomes. By contrast, pronounced diversity effects emerged in the virus-mediated multistrain model (Fig. 4a). Increasing number of strains substantially altered invasion probabilities compared to the coarse-grained outcomes. We quantified the critical viral adsorption rate 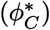 required to reduce invasion probability below 50%. Greater phage diversity enhanced community resilience by decreasing critical viral adsorption rates. In particular, at *γ*_*CB*_ = 0, increasing diversity from *n* = 1 to *n* = 30 reduces the critical adsorption threshold by approximately 60%. These results indicate that phage diversity strengthens the ecological firewall against invasion by expanding the parameter space associated with pathogen exclusion.

**Figure 4.**
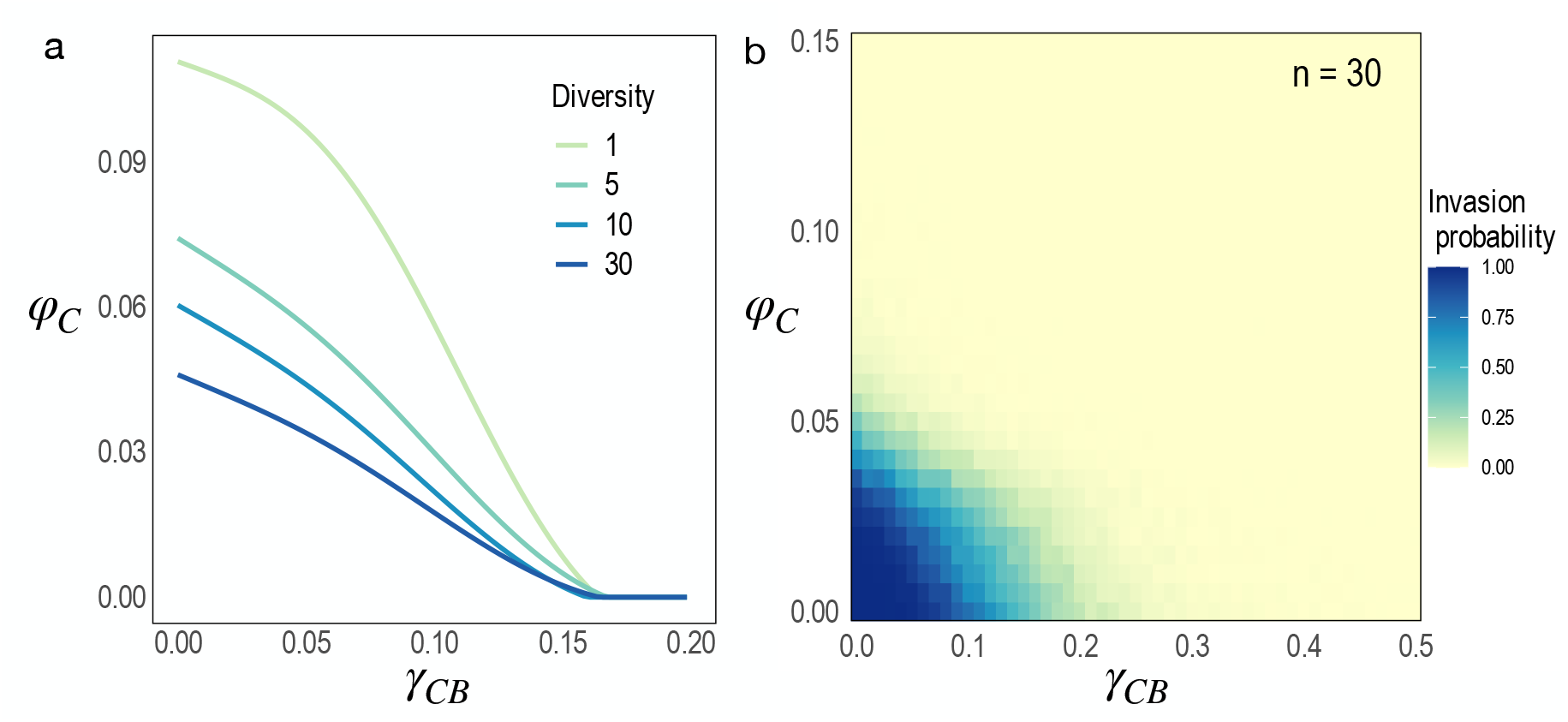
Diversity strengthens pathogen exclusion. (a) Critical viral adsorption rate 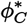 required to reduce invasion probability below 50% as a function of bacterial suppression strength *γ*_*CB*_ . Increasing phage diversity progressively lowers the adsorption threshold required for pathogen exclusion, expanding the parameter space in which pathogens fail to establish. (b) Invasion probability for a representative highly diverse virome (n=30). Darker colors indicate a greater probability of successful pathogen establishment.

## IV. DISCUSSION

Our results strongly support that bacteriophages act as a top-down ecological control and as a key component of microbiome recovery and pathogen clearance. Viral control has long been recognised as a crucial regulator in nutrient cycling and microbial diversity [3, 26, 50] and most recently in antibiotic-induced dysbiosis treatments [38]. We showed that pathogenic invasion depends not only on average viral adsorption or bacterial suppression rates, but on initial microbiome states as well. Such multistability in gut systems is increasingly recognized as a common feature of microbial ecosystems, where competitive interactions and nonlinear feedbacks generate multiple stable community configurations under identical environmental conditions [2, 9, 16, 44]. This phenomenon is the simplest form of memory. Such history-dependent dynamics are common in ecological systems, where priority effects and nonlinear interactions allow past states to influence future community trajectories [11, 17]. Alternative stable states provide a mechanistic explanation for inconsistent clinical outcomes after viral transplantation. Furthermore, we identified a specific state-space region of recurrence susceptibility, where a small perturbation in pathogen or microbial density can switch between pathogen clearance or persistence. This separatrix between basin boundaries implies high uncertainty in microbiome recovery which is not only history-dependent but sensitive to perturbations. These results align well with the reported empirical relapse rates after microbiome transplantations, typically around 20% in clinical studies [23].

We clearly showed that virome reshapes invasion thresholds. Our results distinguish two qualitatively different modes of pathogen control; competition and phage threshold. Bacterial suppression reduces pathogen density gradually through competitive effects, whereas phage adsorption introduces a hard threshold, whereby once adsorption exceeds a critical value, pathogen clearance becomes inevitable. This nonlinear effect generates a regime shift between dysbiotic and healthy states, providing a mechanistic basis for abrupt recovery or relapse. An interesting implication of our results is that path dependence in the system is governed by the initial balance between the resident microbiome and the pathogen rather than by the initial viral population. This implies that virome-based interventions may benefit more from phages with strong adsorption efficiency than from increasing the volume or diversity of virome transplants, potentially allowing for more universal treatment strategies.

Our model introduces a “firewall” mechanism [34, 48]. If the virome is maintained at a positive equilibrium *P*^∗^ *>* 0 by the resident microbiome, then phage predation provides an additional, density-independent removal pressure on the pathogen at the moment of invasion. In effect, the resident microbiome supports a standing virome that can immediately reduce *C*.*difficile* fitness upon introduction. The system thus exhibits an emergent community-level defense [5]: the microbiome indirectly protects itself by sustaining a virome that selectively targets the pathogen while tolerating resident bacteria. Increasing *P*^∗^ strengthens the firewall through *ϕ*_*C*_*P*^∗^, but phage predation also imposes a cost on the resident microbiome through the term *ϕ*_*B*_*PB*. Thus, a protective virome is most effective when it combines high pathogen targeting (*ϕ*_*C*_) with relatively modest resident targeting (*ϕ*_*B*_), or when the resident microbiome compensates via higher net growth *r*_*B*_ and positive internal interactions *α*. This defense mechanism is amplified with increasing diversity. In multi-strain communities, the top-down phage control increases the likelihood of pathogen clearance, emphasizing the stabilizing role of complex virome-microbiome interactions.

Our model assumes mean-field interactions without spatial structure. Moreover, parameters are not calibrated to specific experimental systems. Targeted *in vitro* experiments can be used to test eradication thresholds and recurrence susceptibility. These simplifications allow analytical tractability but limit biological realism. Future work can be extended to incorporating spatial structure, migration, and network structure, which would further clarify how gut virome shapes stability landscapes. Overall, our results suggest that the virome acts as key regulators of resilience, invasion resistance, and recovery in gut microbial ecosystems. The ecological principles identified here may extend beyond the gut microbiome. Virome-mediated control could inform intervention strategies in soil or aquatic microbiomes.

## V. MATERIALS AND METHODS

### A. Microbiome–phage–pathogen model of post-FVT dynamics

We developed a deterministic dynamical model describing interactions between commensal bacteria, bacteriophages, and a pathogenic population representing Clostridioides difficile. The system includes three interacting compartments: a resident microbiome composed of bacterial populations {*B*_*i*_} a virome composed of bacteriophage populations {*P*_*j*_}, and a pathogen population *C*. Population dynamics are governed by a generalized Lotka–Volterra framework that incorporates logistic growth, interspecific microbial interactions, and phage adsorption dynamics. The full model consists of *N* + *M* + 1 coupled ordinary differential equations describing the abundances of bacterial strains *B*_*i*_, phages *P*_*j*_, and the pathogen *C*. The resulting system captures the combined effects of bottom–up resource limitation, microbial competition, and top–down viral predation in shaping microbiome stability and pathogen invasion dynamics. The coarse-grained Bacteria-C. difficile model reads as:

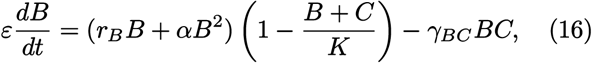

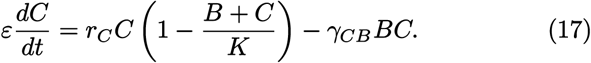

Here, the effective rates growth *r*_*B*_, *r*_*C*_, interaction coefficient *α*, and bacterial suppression rates *γ*_*BC*_, *γ*_*CB*_ are obtained by averaging over the original species-specific rates (see SM). The virus-mediated model reads as:

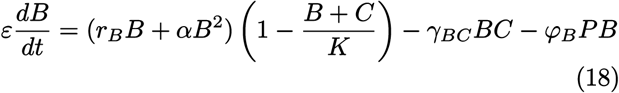

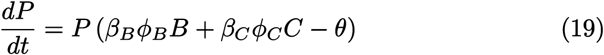

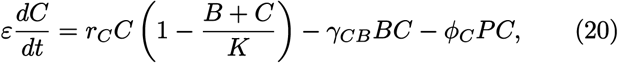

where the bacteriophages *P* exert top-down control of *C. difficile* and Bacteria with the effective adsorption rates *ϕ*_*C*_ and *ϕ*_*B*_ together with the effective burst rates of *β*_*C*_ ans *β*_*B*_. The full model formulation and parameter definitions are provided in the Supplementary Materials.

### B. Full dynamical model of antibiotic exposure and FVT intervention

The analytical results presented above focus on the post-treatment ecological dynamics after FVT has altered the virome composition. For analytical tractability, the virome is therefore represented by a single effective phage compartment *P* . To explicitly model the sequence of antibiotic treatment, dysbiosis, FVT administration, and recovery, we extend the model below by separating the virome into resident phages (*P*_*R*_) and transplanted phages (*P*_*F*_ ). This distinction is used only in the intervention model and does not affect the analytical results derived for the post-treatment state.

We investigated the ecological dynamics starting from the antibiotic exposure to the virome transplantation. We developed a coarse-grained dynamical model consisting of four interacting compartments: resident commensal bacteria (B), resident phages (*P*_*R*_), transplanted FVT phages (*P*_*F*_ ), and the pathogen C.difficile (C). The model incorporates bacterial growth, microbiome self-facilitation, phage predation, pathogen suppression, antibiotic perturbation, and phage transplantation. Antibiotic treatment is represented as a temporary increase in resident bacterial mortality. During the treatment window [*t*_*AB*_,*t*_*AB,end*_), commensal bacteria experience an additional mortality rate *µ*_*AB*_, reducing microbiome abundance and weakening colonization resistance. FVT is subsequently administered at time *t*_*F V T*_ as a pulse inoculation of transplanted phages, increasing *P*_*F*_ by an amount *I*_*F V T*_ . The separation of resident and transplanted phages allows FVT efficacy to emerge mechanistically from differences in adsorption rates (*ϕ*), burst sizes (*β*), and host specificity, rather than being imposed through ad hoc changes in pathogen growth or mortality parameters.

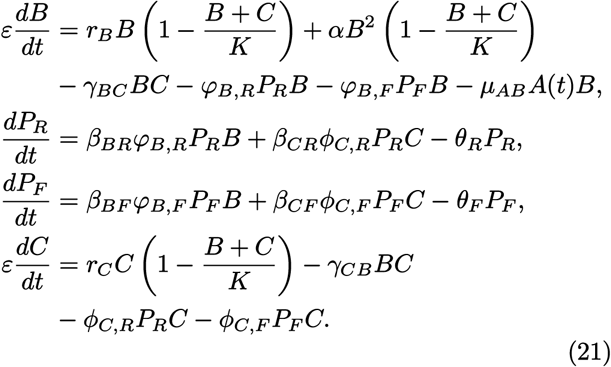

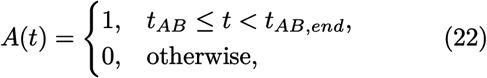

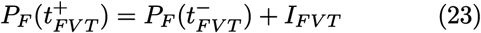

### C. Invasion probability and recurrence susceptibility

In our model, the resident microbiome and the pathogen compete for a shared resource (carrying capacity, *K*), which introduces nonlinear feedbacks between bacterial and pathogen populations leading to multiple stable states. In particular, the system admits two alternative attractors corresponding to pathogen clearance (*C*^∗^ = 0) and pathogen persistence (*C*^∗^ *>* 0), which depend on the initial conditions of the system leading to path-dependence. To quantify how pathogen invasion depends on initial microbiome states, we analyzed the *basins of attraction* of the coarse-grained models. For each parameter combination, we generated *N* = 10^4^ random initial conditions for the resident microbiome (*B*_0_), pathogen (*C*_0_), and phage (*P*_0_) densities by sampling uniformly within biologically feasible bounds: *B*_0_, *C*_0_, *P*_0_ ∼ *U* (0, *K*). The system of ordinary differential equations was numerically integrated until convergence (*B*^∗^, *C*^∗^, *P*^∗^). Each trajectory was classified according to the long-term pathogen outcome: clearance if *C*^∗^ = 0 and persistence if *C*^∗^ *>* 0.

The *invasion probability* was quantified as the fraction of sampled initial conditions that converge to pathogen persistence. Then, we quantified *recurrence susceptibility* of the system as the area of state space lying near the basin boundary separating pathogen clearance from persistence. This region represents microbiome states for which small perturbations in initial bacterial or pathogen densities can change the long-term outcome. Let *I*_clear_ and *I*_persist_ denote the sampled initial conditions leading to pathogen clearance and pathogen persistence, respectively. Because these two sets are disjoint by definition, recurrence susceptibility was estimated from the overlap between their geometric envelopes in the (*B*_0_, *C*_0_) state space. We define ℋ_clear_ and ℋ_persist_ as the concave hulls enclosing *I*_clear_ and *I*_persist_. Recurrence susceptibility is then given by

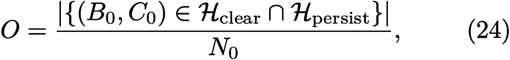

Here, | · | denotes the area in the (*B*_0_, *C*_0_) plane. Large values of *O* indicate that the clearance and persistence basins are geometrically intermixed, whereas *O* = 0 indicates well-separated basins. In other words, the recurrence-susceptibility region represents the set of microbiome states from which both pathogen clearance and persistence are possible depending on small perturbations (see more details in SM).

### D. Multi-strain simulations

To investigate the effect of microbial diversity on invasion dynamics, we extended the coarse-grained models to include multiple bacterial and viral strains. Communities were simulated with varying numbers of strains *n* = {5, 10, 30}, where *n* represents the number of bacterial and viral strains in the system. Interaction coefficients and growth rates were sampled from lognormal distributions centered around the coarse-grained parameter values to capture ecological variability among taxa. A sparse host-range matrix was imposed on phage-host interactions to represent limited viral host specificity. Simulations of these multi-strain systems were used to evaluate how increasing bacterial and viral diversity modifies invasion probability. Full details of the multi-strain parameterization and sampling procedures are provided in the Supplementary Materials.

## ACKNOWLEDGMENTS

The authors thank the members of the Complex Systems Lab for valuable discussions and feedback. RS acknowledges the support of the AGAUR 2021 SGR 0075 grant, a AEI-PID2023-152129NB-I00 grant, and by the Santa Fe Institute.

## Notes

### Competing Interest Statement

The authors have declared no competing interest.

### Summary of Updates

This revised version includes expanded analytical results on pathogen eradication thresholds, microbiome recovery dynamics, and diversity effects. Several figures have been redesigned, and the Introduction, Results, Discussion, Methods have been extensively revised to improve clarity and presentation.

